# Light diffusion in layered media: A numerical study in the spatial and time-domains

**DOI:** 10.1101/2022.04.26.489577

**Authors:** Michael Helton, Samantha Zerafa, Karthik Vishwanath, Mary-Ann Mycek

## Abstract

Accurate and efficient forward models of photon migration in heterogeneous geometries are important for many applications of light in medicine because many biological tissues exhibit a layered structure, with each layer having independent optical properties and thickness. Unfortunately, closed form analytical solutions are not readily available for layered tissue-models, and often are modeled using computationally expensive numerical techniques or theoretical approximations that limit accuracy and real-time analysis. Here, we develop an open-source accurate, efficient, and stable numerical routine to solve the diffusion equation in the steady-state and time-domain for a layered cylinder tissue model with an arbitrary number of layers and specified thickness and optical coefficients. We show that the steady-state (< 0.1 ms) and time-domain (< 0.5 ms) fluence (for an 8-layer medium) can be calculated with absolute numerical errors approaching machine precision. The numerical implementation increased computation speed by 3 to 4 orders of magnitude compared to previously reported theoretical solutions in layered media. We verify our solutions asymptotically to homogeneous tissue geometries using closed form analytical solutions to assess convergence and numerical accuracy. Approximate solutions to compute the reflected intensity are presented which can decrease the computation time by an additional 2-3 orders of magnitude. We also compare our solutions for 2, 3, and 5 layered media to gold-standard Monte Carlo simulations in layered tissue models of high interest in biomedical optics (e.g. skin/fat/muscle and brain). The presented routine could enable more robust real-time data analysis tools in heterogeneous tissues that are important in many clinical applications such as functional brain imaging and diffuse optical spectroscopy.

## Introduction

Optical properties can be used as indicators of pathological and physiological conditions of biological tissue^1,2^. Accurate quantitation of these properties from experimental measurements depend on analytical models that need to account for the structural complexity of the tissue system. Therefore, it is important to consider the optical heterogeneity of biological tissues, which are usually approximated as optically homogeneous to facilitate data analysis^3,4^. Experimentally, light propagation measurements are made by illuminating the tissue surface with either a continuous, frequency modulated, or pulsed light source and collecting measurements of the scattered light after it has propagated through the tissue medium^5,6^. Measured optical signals are translated into absorption and scattering properties of the medium by utilizing an appropriate forward model of light transport that best represents the measured data^7,8^.

Light propagation in random media such as biological tissues is theoretically modeled using the Radiative Transfer Equation (RTE)^9–11^. Due to the highly scattering nature of these media, the RTE can be reduced to the diffusion equation, which gives analytical solutions in homogeneous, semi-infinite, or infinite slab geometries^12,13^. The RTE can also be solved by the Monte Carlo method which remains the gold-standard approach to calculate light transport in media with complex geometries^14^ but is computationally expensive^15^. Although parallel implementations have significantly improved the speed of Monte Carlo simulations^16,17^, they still broadly remain non-viable as inverse solvers to obtain optical properties from experimental measurements^16^.

Theoretical approaches that account for structural complexity in tissues provide improved reconstruction of optical properties using diffuse optical measurements when studying brain hemodynamics^18^. Although Monte Carlo methods can simulate light propagation in realistic head geometries derived from magnetic resonance imaging (MRI) data^16^, modeling the head as layered homogeneous slabs, each with their own set of optical properties, provided similar accuracy in reconstruction of optical properties^3^. Further, diffuse optical measurements are applicable to various parts of the body that exhibit a layered structure (e.g. skin over top muscle, scalp and skull surrounding brain tissue). Therefore, an accurate, versatile and efficient analytical approach to model spatially and/or temporally resolved diffuse reflectance in layered media would enhance optical property reconstructions from diffuse optical measurements obtained in such layered media *in vivo*.^3^.

Several methods to solve the diffusion equation for layered media have been reported in literature by using integral transforms^19–22^, method of images^23^, eigenfunctions^24,25^, or finite differences^26^. These methods do not give closed-form expressions directly in the spatial or time-domains for the photon fluence. Instead, the fluence in real-space is computed using numerical transforms^22^ or root-finding techniques^24^ which tend to increase numerical errors and computational costs^27^. For example, the integral transform approach^22^ solves the diffusion equation in the spatial-frequency domain which then must be inverse space-transformed (e.g. 2-D inverse Fourier) for real-space calculations. An additional inverse time Fourier transform is required for computation in the time-domain^22^. Both of these transforms make calculations of the steady-state and time-domain solutions difficult to compute for a wide range of optical and geometrical inputs^19,22^. Other approaches have been developed to compute geometries with large layer thicknesses and high scattering coefficients and/or spatial frequencies but rely on approximations^19,28–30^. Given these challenges, the fastest reported computational times for time-domain fluence in multi-layered media range ≈0.5-5 seconds, depending on the number of layers and numerical accuracy required^21,24,28,31^. Such computational performance would preclude direct use of such layered analytical solutions for real-time analysis as optimization of multiple parameters in layered media would take several minutes^32^.

In this report, we present an accurate and efficient procedure for computing the photon fluence in an N-layered cylinder using solutions to the diffusion equation^21^. Our code is open-source and well documented for ease of use. We overcome the computational difficulties noted above by modifying the solutions^21^ in the spatial-frequency domain for numerical stability, which allows for computation of arbitrarily sized inputs without approximations. Lastly, we use an inverse Laplace transform to improve convergence in the time-domain which improved the numerical accuracy while decreasing the computational cost by several orders of magnitude^33,34^. Below we describe: (a) implementation of the numerical solutions in the steady-state and time-domain for diffuse optical reflectance and transmittance measurements in N-layered media, (b) verification of the numerical accuracy and stability of the approach in calculating photon fluence for several source-detector configurations and tissue models, and (c) validation by direct comparisons to Monte Carlo simulations of fluence in multi-layered tissue models.

## Methods

### Theory

In highly scattering turbid media such as biological tissue, the steady-state diffusion equation^35^

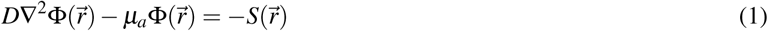

can be used to approximate light propagation in random turbid media where 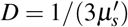, *μ*_*a*_, and 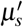 denote the fluence rate, the diffusion coefficient, the absorption coefficient, and the reduced scattering coefficient, respectively^35^. The fluence can be used to calculate the diffusely reflected or transmitted intensity which are the quantities usually experimentally measured^13^. We assume an incident beam can be approximated by an isotropic point source located at a distance of 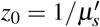 from the incident surface. Using an integral transform, equation 1 can be solved in the spatial-frequency domain under extrapolated boundary conditions considering N-layers of distinct absorption and scattering properties of arbitrary thicknesses^21,35^. For the special case of a point source incident onto the center top of the cylinder, the fluence in the *k*^*th*^ layer can be expressed in real space after applying an inverse finite Hankel transform^21,35^ by

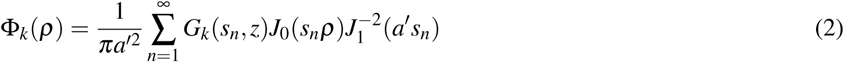

where *J*_*m*_ is the Bessel function of first kind and order *m* and *s*_*n*_ is determined from the roots of *J*_*m*_ such that *J*_*m*_(*a*′*s*_*n*_) = 0, *n* = 1, 2, …, where *n* is the *n*^*th*^ root of *J*_*m*_. *z* is the detector depth within the medium in cylindrical coordinates such that Φ_1_(*z* = 0) is used for reflection calculations and Φ_*N*_(*z* = *L*) where *L* is the sum of all the layer thicknesses is used for transmission. The extrapolated boundary is determined with *a*′ = *a* + *z*_*b*_ where *a* is the radius of the cylinder and *z*_*b*_ = 2*AD* where *A* is proportional to the fraction of photons that are internally reflected at the boundary^12^. *G*_*k*_ represents the Green’s function in the *k*^*th*^ layer which must be solved separately for each layer. Derivations of equation 2 are given in Appendix A of the Supplementary material along with the explicit expressions of *G*_*k*_ in the topmost and bottommost layers. Applying inverse transforms directly to *G*_*k*_ as previously given^21,35^ can lead to numerical overflow for large layer thicknesses, scattering coefficients, and frequencies. To overcome these issues, we give expressions for *G*_*k*_ in terms of exponentially decaying terms that will not cause numerical overflow for large arguments and do not need to make any approximations.

Time-domain solutions are often computed using the inverse Fourier transform by using the substitution *µ*_*a*_ → *µ*_*a*_ + *iω*/*c* and computing the real and imaginary parts of the fluence in the frequency domain at many frequencies (400-4,000)^21,33^. As these Fourier integrals are slow to converge and can rapidly oscillate, the number of nodes needed for Gaussian integration methods are highly dependent on input parameters which makes the Fourier integral difficult to accurately and efficiently compute^33^. Instead, an inverse Laplace transform with 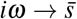 can be used and numerically integrated along a complex contour^33,34^. Utilizing a contour that begins and ends in the left hand plane Re *z* → −∞ forces a rapid decay of the integrand making for easier and faster numerical integration by using trapezoidal rules^34^. The corresponding solution for the time-domain fluence is then

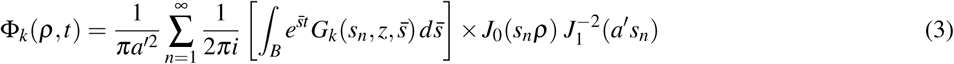

where *B* denotes the Bromwich path with 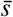 being a complex number along the contour. A single contour is considered for rapid evaluation of the fluence at many time points for *t* ∈ (*t*_1_, *t*_2_)^33,34^. The full derivation of the time-domain solution shown in equation (3) along with details of the numerical Laplace transform are provided in Appendix B of the supplementary material.

### Numerical Verification

To calculate the steady-state fluence in real space, a finite inverse Hankel transform (equation (2)) must be numerically computed. While calculation of solutions in the time-domain requires equation (2) to be evaluated for *N* complex valued absorption terms during the numerical inversion of the Laplace transform in equation (3). The numerical accuracy and efficiency of the procedure depend on the convergence and difficulty of computing the two sums. Since both the computation of the steady-state and time-domain fluence depends on equation (2), the accuracy and computational speed depend primarily on how many terms *n* of the infinite sum are retained in equation (2).

As exact, closed-form solutions for the photon fluence in layered media are not available, we first validate our solutions in layered media to closed-form homogeneous solutions for semi-infinite media^13^. In these validations, each layer in our layered tissue-model was set to have the same optical properties as the homogeneous medium along with laterally infinite boundaries. This allows us to precisely quantify numerical errors and determine convergence of our solutions in terms of the number of terms retained in the sum in equation (2). We first compare equation (2) evaluated with 2 and 8 layers of similar optical properties to the semi-infinite solution^13^ and to Monte Carlo simulations in a semi-infinite medium. Next, we compare the absolute accuracy of the routine as a function of *n* compared to the semi-infinite solution to assess the rate of convergence of equation (2) considering different input parameters 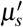, *µ*_*a*_, *z* and *a*.

Solutions in the time-domain require computing both the infinite sum in equation (2) as well as numerically performing the inverse Laplace transform in equation (3). Strategies to invert the Laplace transform could allow for significantly faster convergence compared to the Fourier transform^33,36^, however the obtention of the inverse Laplace transform is not always an easy or even possible task to perform^36^. Therefore, the accuracy of the numerical approach to invert the Laplace transform must be rigorously tested. We focus on two attributes in performing the numerical integration in equation (3) that affect both the convergence and numerical accuracy: (a) the number of Laplace space evaluations *N* used to evaluate the Laplace integral in equation (3) and (b) the contour width determined from Λ = *t*_2_/*t*_1_ where *t* ∈ (*t*_1_, *t*_2_). The effect of both of these parameters on the convergence and numerical accuracy are again examined by comparison to closed-form homogeneous analytical solutions. We show the reconstruction of the time-domain signal for high scattering and high absorbing media at short and long distances and times where the numerical reconstruction has been difficult to perform^30,33^. We have included in Appendix A extended discussion on how to efficiently compute equation (2), which also directly affect the computation of equation (3), and the advantages compared to other routines^19^. Additionally, we give approximations for reflectance simulations (*z* = 0) that are accurate for double precision arithmetic (see Supplementary Fig. S3) which can decrease the computational time by 2-3 orders of magnitude.

All the numerical routines and figures presented here were developed using the Julia programming language (v1.7.0)^37^. Numerical simulations were performed on a MacBook Pro with an Apple M1 chip (MacOS version 11.1) and 16 GB of memory. Simulations in the steady-state utilized a single core while the inverse Laplace transform in the time-domain used multi-threaded parallelism. Here, the Laplace space evaluations were evenly distributed across the 4 cores and 8 threads of the M1 chip. All benchmarks are done in double precision arithmetic using v0.8.0 of LightPropagation.jl.

### Validation with Monte Carlo

To validate the derived analytical solutions, the fluence is compared with results obtained from Monte Carlo simulations. The Monte Carlo method simulates the propagation of photons through the scattering medium using appropriate probability functions and random number generation^15,16^. In the limit of an infinitely large number of photons used in the simulations, the Monte Carlo method is an exact solution of the RTE^15^. We utilized an independent open-source Monte Carlo code provided by the Virtual Photonics Technology Initiative^38^ to validate the layered diffusion theory model for three biologically relevant tissue models. The Monte Carlo simulations used 5 × 10^7^ photons for each simulation with an isotropic emitting source at a depth of 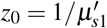 and a Henyey-Greenstein phase function. The anisotropic factor was assumed to be *g* = 0.8 for all layers. For all comparisons the refractive index of the medium is assumed to be *n* = 1.4 where the external medium is assumed to be air *n* = 1.0. The fluence as a function of *t* and/or *ρ* and *z* was recorded in discrete bin widths of Δ*t* = 0.02 ns, Δ*ρ* = 0.99 mm, and Δ*z* = 0.27 mm.

## Results

### Numerical accuracy of the layered diffusion equation

In Fig. 1, the fluence on the top boundary (*z* = 0) in a semi-infinite medium with optical coefficients *µ*_*a*_ = 0.1 cm^−1^, 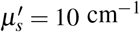, *g* = 0.8, *n* = 1.4 is simulated using Monte Carlo methods in the steady-state and time-domain. We compare the results to solutions of the diffusion equation in a semi-infinite medium^13^ and to equations (2) and (3) when solved for a 2 and 8 layered medium with the same optical coefficients in each layer. Here, we used a cylinder radius of *a* = 20 cm and a total cylinder length *L* of 10 cm (i.e., the thickness of each layer in the 2 and 8 layered model was 5 and 1.25 cm, respectively) to approximate a semi-infinite medium. As previously shown^13^, diffusion theory exhibits excellent agreement with relative errors (|1−Φ_*DT*_ /Φ_*MC*_|) < 0.05 compared to Monte Carlo simulations given enough scattering events. Equation (2) also shows excellent agreement to the closed-form semi-infinite solution^13^ in both the steady-state and time-domain giving similar relative errors to the Monte Carlo results.

**Figure 1.**
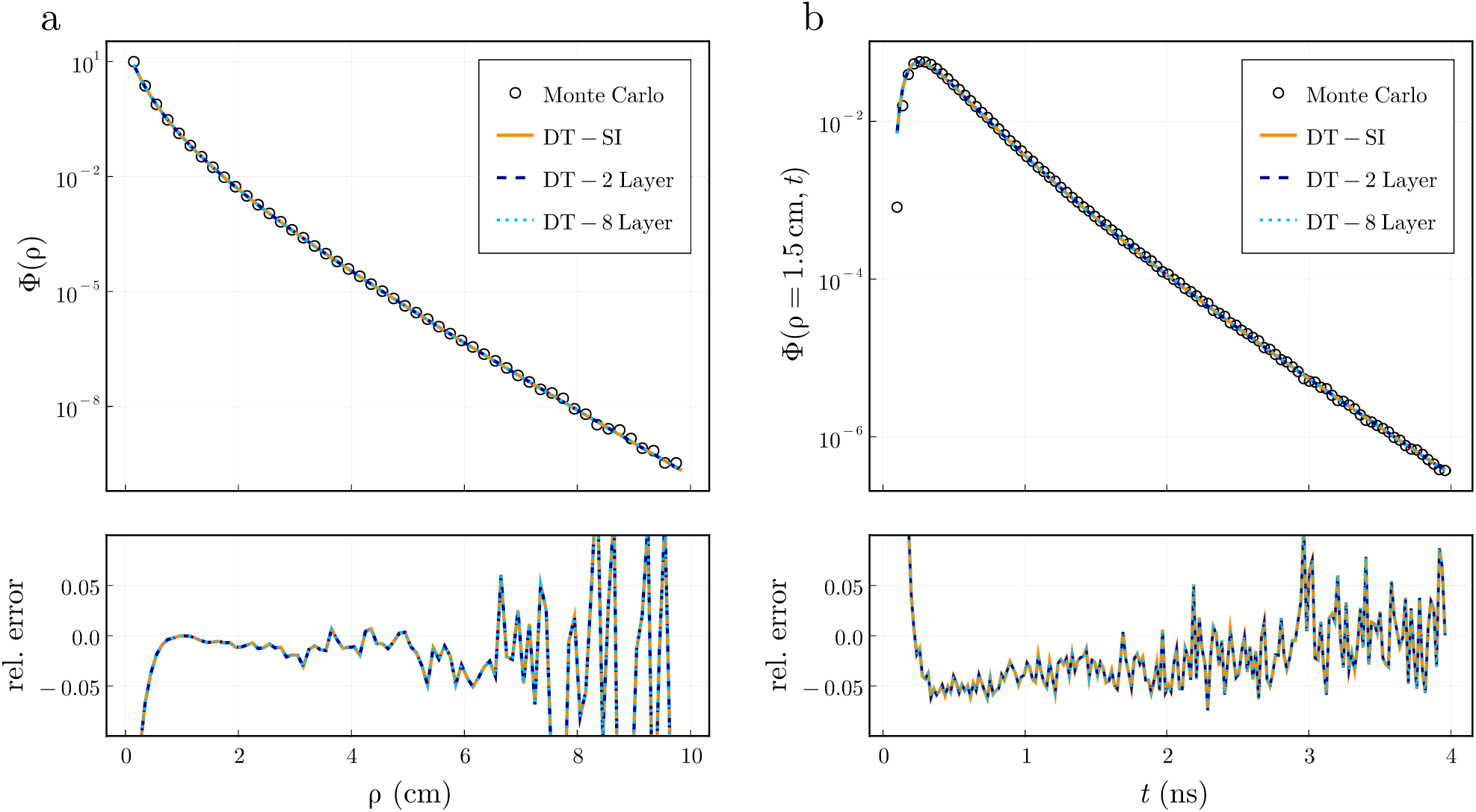
Equations (2) and (3) computed for both 2 and 8 layers agree with Monte Carlo simulations within relative errors of 0.05 which matches the errors achieved with the semi-infinite (SI) solution^13^. We show the (a) steady-state and (b) time-resolved fluence calculated with Monte Carlo simulations and diffusion theory for a semi-infinite medium with optical coefficients *µ*_*a*_ = 0.1 cm^−1^, 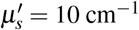, *g* = 0.8, *n* = 1.4 and *z* = 0 cm. We considered the same optical properties in each layer and laterally infinite geometries in (2) and (3) to approximate semi-infinite media. The relative error between the diffusion theory results and Monte Carlo are shown below.

In contrast to the semi-infinite solution^13^, the numerical accuracy of equation (2) is affected by the termination of an infinite sum after *n* terms. For example, given a large number (≈1, 500) of *n*, the layered simulations shown in Fig. 1 can approximate the closed-form semi-infinite solution close to the limits of the numerical precision (detailed below). However, the sum is usually terminated after a desired tolerance is reached as was done in Fig. 1. In Fig. 1a, the steady-state fluence used *n* = 500 for *ρ* < 2 cm, *n* = 1000 for *ρ* < 7 cm, and *n* = 1500 for *ρ* < 10 cm whereas in Fig. 1b we use just *n* = 50 for both the 2 and 8 layer simulations. In general, to simulate lower fluence values a larger number of roots in equation (2) will be required to achieve similar relative errors. Consequently, the number of terms *n* required in equation (2) will be dependent to varying degrees on the input optical properties and cylinder dimensions considered.

In Fig. 2, we investigate how the convergence of equation (2) is dependent on the input parameters 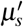, *µ*_*a*_, *z*, and *a* by showing the absolute error (|Φ_*SI*_−Φ_*Lay*_|) between the layered solution (Φ_*Lay*_) simulated with equation (2) and the semi-infinite solution (Φ_*SI*_)^13^. We use baseline optical properties for a 2 layer medium of 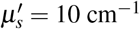, *µ*_*a*_ = 0.1 cm^−1^, *l* = (1.0, 20.0) cm, *a* = 10 cm, and *ρ* = 1 cm, where the optical properties are the same in both layers to approximate a semi-infinite medium. Although we use the same 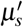 in both layers, the convergence of equation (2) is highly dependent on the scattering coefficient in the first layer 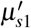 as seen in Fig. 2a. Increasing 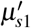 severely diminishes the convergence of equation (2) when *z* = 0 (see Supplementary material in Appendix A for extended discussion). On the other hand, for the range of values shown here, *µ*_*a*_ had a negligible effect on the convergence (Fig. 2b). There is a close relationship between 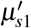 and *z* as shown in Fig. 2a and 2c and their effect on the convergence of equation (2). When *z* ≈ *z*_0_ with 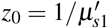, equation (2) requires a high number of terms to converge. This is also the primary reason why increasing the scattering coefficient also requires significantly more terms when *z* = 0 as *z*_0_ ≈ *z*. Additionally, increasing *a* results in slower convergence due to smaller values of *s*_*n*_ during the sum. We note that for the situations where 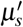, *µ*_*a*_, *z*, and *a* are small, the semi-infinite approximation is less valid resulting in worse agreement between the semi-infinite solution even though the sum quickly converges. The routine can be made accurate down to absolute errors of the machine precision used in the calculation. For example, Fig. 2 was calculated using double precision arithmetic with machine precision *ε* ≈ 10^−16^. The loss of precision in calculating *J*_0_(*s*_*n*_*ρ*) in equation (2) is the primary limitation of the routine. For typical 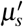 found in biological tissue 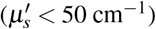, less than 1,000 roots are usually sufficient. For example, only 50 terms were used in Fig. 1b resulting in similar relative errors compared to Monte Carlo simulations when using 5,000 terms.

**Figure 2.**
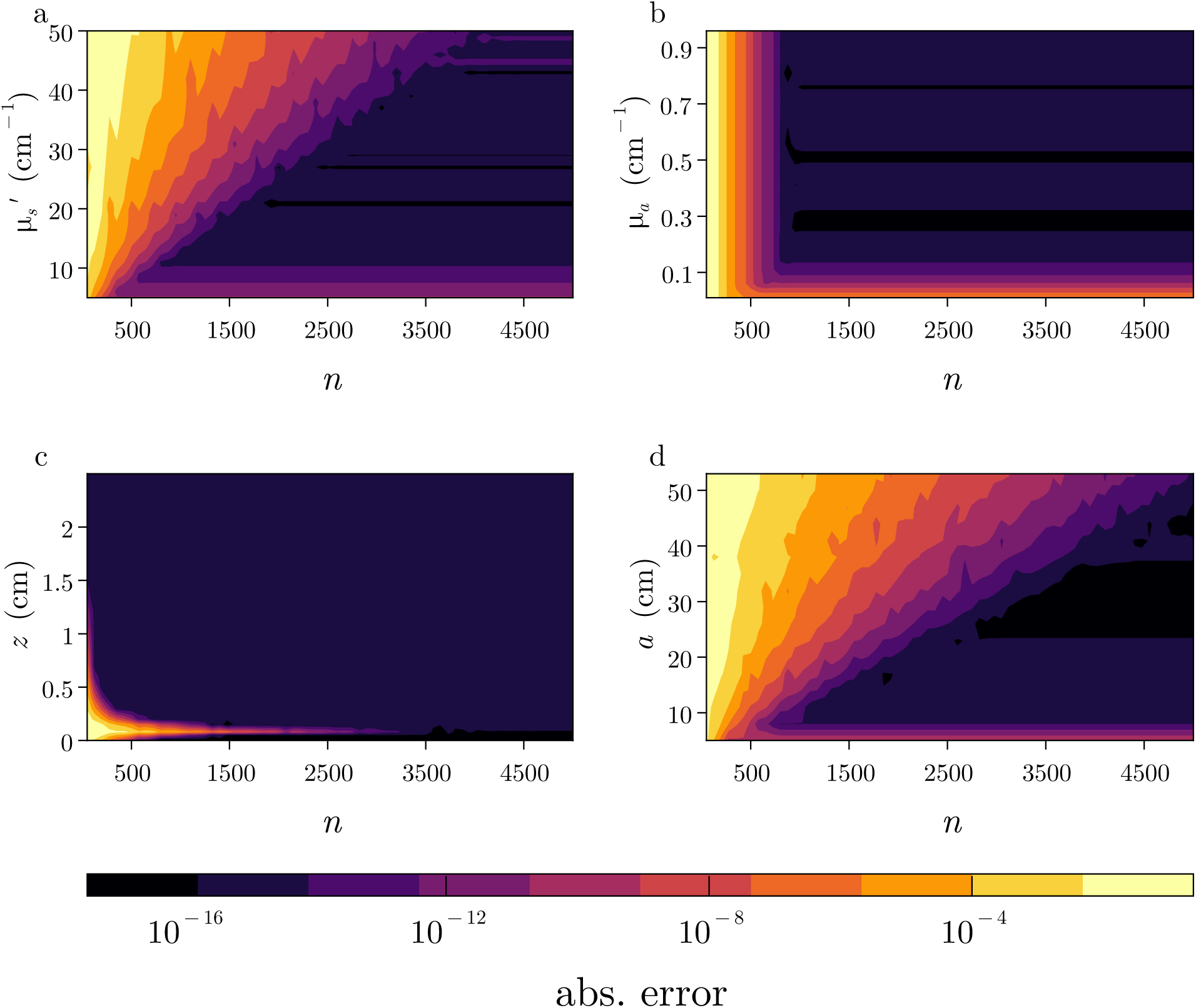
The rate of convergence of the infinite sum in equation (2) depends mostly on input parameters 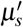, *z*, and *a* while showing little dependence on *µ*_*a*_. We show the absolute error between the semi-infinite solution^13^ and equation (2) as function of the number of terms *n* used in (2) for different values of (a) 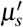, (b) *µ*_*a*_, (c) *z*, and (d) *a*. We fix the other properties to 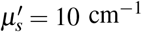, *µ*_*a*_ = 0.1 cm^−1^, *l* = (1.0, 20.0) cm, *a* = 10 cm, and *ρ* = 1 cm while using the same optical properties in each layer.

The accuracy of the time-domain solution given in equation (3) is affected by both the termination of the sum in equation (2) as previously discussed and the numerical inversion of the Laplace integral in equation (3). We focus on two main attributes for the convergence of the inverse Laplace transform: (a) the hyperbola contour size (proportional to Λ = *t*_2_/*t*_1_) and (b) the number of Laplace space evaluations *N* used to evaluate the Laplace integral in equation (3) by comparing the time-resolved fluence simulated with equation (3) to the semi-infinite solution^13^. The fluence is simulated at *ρ* = 1 cm on the top boundary (*z* = 0) using a 4 layered model with layer thicknesses of *l*_*k*_ = (0.5, 1.5, 3.0, 5.0) cm, 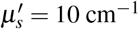 and *µ*_*a*_ = 0.1 cm^−1^ with a radius of 15 cm to approximate a semi-infinite geometry. Typically, the time-domain signal is required at many values in some range *t* ∈ (*t*_1_, *t*_2_) where it becomes significantly more efficient to use a single contour for all time points^33^.

In Fig. 3a, we show the absolute (top) and relative (bottom) errors between equation (3) and the semi-infinite solution^13^ at a single instant of time *t* = 1.0 ns as a function of *N*, for four different values of Λ. Variable values of Λ are achieved by using different *t*_1_ values of 1.0, 0.1, 0.01, and 0.001 ns such that Λ*t*_1_ = 5 ns is fixed and *t* = 1 ns is within the bounds of (*t*_1_, *t*_2_). The absolute errors were similar for any *t* value within *t* ∈ (*t*_1_, *t*_2_) while the relative error was dependent on the value of *t* (i.e. larger relative errors are observed at long times when the fluence is lowest). Less than 20 Laplace evaluations were needed to give absolute errors < 10^−8^ even for large values of Λ. The lowest absolute error was again limited by the machine precision in the numerical procedure.

**Figure 3.**
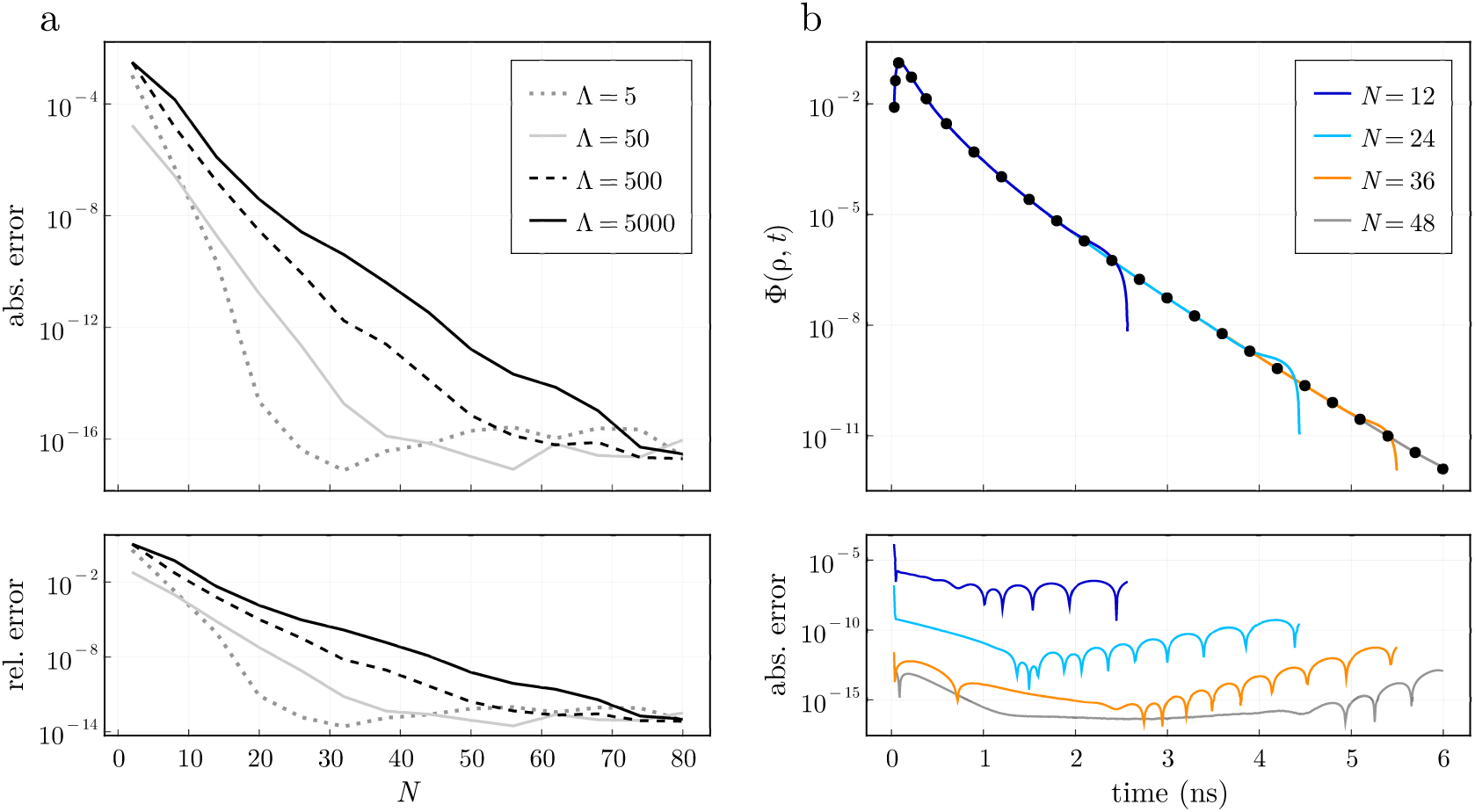
(a) The (top) absolute and (bottom) relative errors for the time-domain reconstruction at a single time value *t* = 1 ns in (*t*_1_, Λ*t*_1_) between the time-domain solution in equation (3) and the semi-infinite solution as a function of the number of Laplace space evaluations *N*. Larger contour sizes (∝ Λ) require higher values of *N* to reach similar accuracies. (b) Reconstruction of the time-domain signal at 600 time points in *t* ∈ (0.03, 6.0) corresponding to Λ = 200 considering four different values of *N*. The semi-infinite solution is shown as black circles with the resulting absolute error between the semi-infinite and layered solution shown in the bottom plot. The absolute error is dependent on *N*, which is similar for all time values considered in *t* ∈ (*t*_1_, *t*_2_)

In Fig. 3b, we considered a single contour Λ = 200 to reconstruct 600 time points in *t*∈ (0.03, 6.0) and we show these for four different values of *N*. A larger *N* improved the overall accuracy and was relatively independent of the time point *t* ∈ (0.03, 6.0) for a given *N*. For larger contours Λ = *t*_2_/*t*_1_ a higher number of *N* are needed to reconstruct the time-domain signal over the whole time window *t* ∈ (*t*_1_, *t*_2_) for a given absolute error. For example, Fig. 3b shows the same Λ but reconstructs the time-domain signal for different values of *N*. However, smaller values of *N* are not able to reconstruct accurately over the entire time window due to the lower fluence values Φ(*ρ, t*) at later times. A given *N* reconstructs the time-domain signal over the entire window at a relatively fixed absolute error. Therefore, larger relative errors will be observed at later times when the fluence is lowest.

We note that calculations in Fig. 3b are only shown up to the point where the time-domain signal is not accurately reconstructed. Even for a rather large value of Λ = 200, only 12 Laplace evaluations were needed to reconstruct the time-domain signal with a dynamic range of 3 orders of magnitude and with 24 evaluations that increased to 6 orders of magnitude, and represent typical scale of dynamic range reported for time-domain systems^8,39^. Increasing the number of evaluations does not decrease absolute errors once the errors reach the machine precision. Coincidentally, we have found that the numerical inversion of the Laplace transform is also limited by absolute errors approaching the machine precision, similar to the numerical computation in the spatial domain.

The previous examples have focused on modest values of 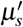, *µ*_*a*_, *ρ*, and layer thicknesses. In Fig. 4, we reconstruct the time-domain signal for high scattering media and large layer thicknesses over a wide range of times which has previously been previously difficult due to numerical overflow^27–29^. In Fig. 4a, the time-domain signal for a high scattering medium 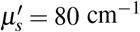 and *µ*_*a*_ = 0.1 cm^−1^ at *ρ* = 0.2 and *ρ* = 3.5 cm is shown. We considered a 4-layered medium with the same optical properties in each layer with layer thicknesses *l*_*k*_ = (0.5, 1.5, 3.5, 30.0) cm and a cylinder radius of 15 cm for comparison to a semi-infinite model^13^. We used *n* = 1, 000 roots in equation (2) with *N* = 24 Laplace evaluations at *ρ* = 3.5 cm and *N* = 72 evaluations at *ρ* = 0.2 cm. Although the fluence at *ρ* = 0.2 cm is significantly larger, we considered *t* ∈ [0.004, 6.0] resulting in a Λ = 1500 whereas at *ρ* = 3.5 we considered *t* ∈ [0.8, 6.0] giving Λ = 7.5. This again highlights that the number of Laplace space evaluations is highly dependent on Λ. Even considering a very large layer thickness *l*_4_ = 30 cm and large reduced scattering coefficient 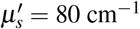, the time-resolved fluence can be easily simulated in double precision arithmetic, very close to the source, and at both early and late times.

**Figure 4.**
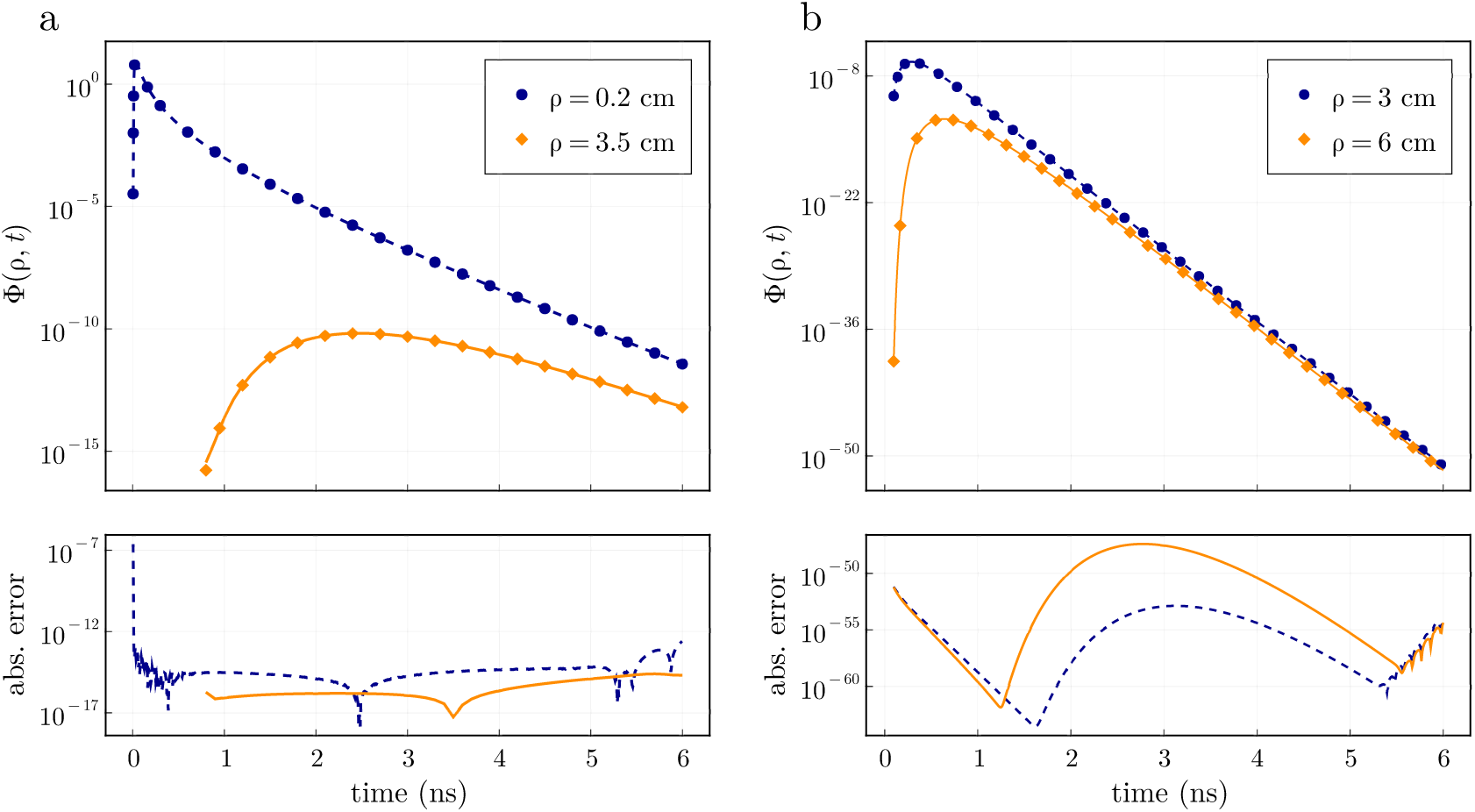
Equation 3 can be computed to absolute errors up to the machine precision compared to homogeneous closed form models at high scattering over a wide range of times and distances away from the source. (a) Time-resolved fluence from a 4-layered highly scattering media with optical properties 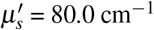 and *µ*_*a*_ = 0.1 cm^−1^ at *ρ* = 0.2 and *ρ* = 3.5 cm. Computation was performed using double precision arithmetic. (b) Time-resolved fluence at the top boundary of a 4-layered media with optical properties 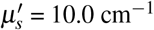 and *µ*_*a*_ = 0.6 cm^−1^ at *ρ* = 3 and *ρ* = 6 cm. Computation was performed using octuple precision arithmetic. The semi-infinite solution is shown as markers with the absolute error between the two solutions shown below.

In Fig. 4b we show calculation of the time-domain fluence at the surface (*z* = 0) for very low fluence values from a high absorption medium (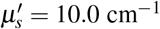 and *µ*_*a*_ = 0.6 cm^−1^) at two detector locations of *ρ* = 3.0 cm and 6.0 cm and for *t* ∈ [0.1, 6.0]. These calculations were performed in octuple precision using *N* = 168 Laplace evaluations and only *n* = 600 roots in equation (2). An increase in amount of Laplace evaluations is needed for very low fluence values and can also be observed by extrapolating the asymptote of convergence in Fig. 3 to very low absolute errors. It is worth noting that it was possible compute time-resolved fluence for over 50 orders of magnitude with high numerical accuracy for source-detector separations as large as 6 cm. Lastly, the roots of *J*_0_ must be calculated in higher precision to achieve the shown absolute errors.

### Comparison to Monte Carlo Simulations

Next, we compared the solutions obtained from equations (2) and (3) to Monte Carlo simulations for both the steady-state and time-domain. We consider three different tissue geometries that are of high clinical interest and have been extensively used to model light propagation in different organ systems previously: a 2-layer model^22^, a 3-layer model representing a skin/fat/muscle layer^30^, and a 5-layer brain model^40^ representing a scalp, skull, cerebrospinal fluid (CSF), and a gray and white matter layer.

The optical properties considered in the 2-layer model are *µ*_*a*1_ = 0.2 cm^−1^, *µ*_*a*2_ = 0.1 cm^−1^, 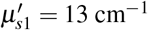, and 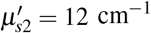 with layer thicknesses of *l*_1_ = 6 mm and *l*_2_ = 90 mm. We report the optical properties of the 3-layer muscle and 5-layer brain model in Table 1. For all tissue models, we consider the index of refraction for each layer to be *n* = 1.4 with the external index of refraction being air (*n* = 1). The anisotropy *g* = 0.8 was consistent for all layers in the Monte Carlo simulations. In all cases we compare the fluence on the top boundary (*z* = 0) as a function of *ρ* for the steady-state calculations and as a function of *t* in the time-domain for *ρ* = 0.45, 1.45, and 3.05 cm.

**Table 1.** Optical properties and layer thicknesses for the 3 layer skin/fat/muscle^30^ and 5 layer brain tissue models^40^Skin, Fat, Muscle Model^30^ Brain Model^40^

| Skin, Fat, Muscle Model<sup>30</sup> |  |  |  | Brain Model<sup>40</sup> |  |  |  |
| --- | --- | --- | --- | --- | --- | --- | --- |
| | $\mu_a \text{ cm}^{-1}$ | $\mu'_s \text{ cm}^{-1}$ | $l \text{ (mm)}$ | | $\mu_a \text{ cm}^{-1}$ | $\mu'_s \text{ cm}^{-1}$ | $l \text{ (mm)}$ |
| skin | 0.15 | 15 | 1.2 | scalp | 0.18 | 19 | 5 |
| fat | 0.02 | 12 | 3.8 | skull | 0.16 | 16 | 8 |
| muscle | 0.2 | 5 | 100 | CSF | 0.04 | 0.25 | 2 |
| - | - | - | - | Gray | 0.36 | 22 | 5 |
| - | - | - | - | White | 0.14 | 9.1 | 40 |

In Fig. 5, we compare the steady-state and time-domain fluence when simulated using the Monte Carlo method and diffusion theory. The left column shows the steady-state fluence Φ(*ρ*) for *ρ* ∈ (0.15, 10) cm simulated using equation (2). Excellent agreement (relative errors < 0.1) is observed for all three tissue models, however the agreement is not uniform. The 2-layer model showed the best agreement for all values of *ρ* where the results asymptotically agreed with the Monte Carlo method. The 3-layer muscle model showed good agreement for *ρ* < 5 cm, but did not asymptotically agree. These results are consistent with recent reports^41^ that showed a breakdown in diffusion theory when the mean free path approaches the thickness of the top layer. Here, a top layer thickness of 1.2 mm was used. Although the significance of these errors were not studied on the reconstruction of optical properties, the relative errors between Monte Carlo solutions are less than 0.1 for *ρ* < 6 cm. Diffuse optical measurements are not usually collected at such large distances due to low signal to noise. A similar effect is observed in the brain model Fig. 5e where agreement (relative error < 0.1) is observed for *ρ* < 6 cm, however longer distances show higher errors. These errors can be mostly attributed to the limitations of diffusion theory to accurately model the low scattering cerebrospinal fluid layer^42^. Our analytical solutions utilized 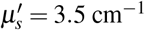 to most accurately model the low scattering CSF layer with diffusion theory as previously suggested^42^, though the choice of 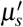 significantly affects the resulting fluence calculated with diffusion theory for *ρ* > 6 cm. If 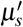 is less than 4 cm^−1^, a severe overestimation of the fluence is seen. Practically, for *ρ* > 6 cm it may become unrealistic to consider the CSF and other brain layers as parallel planes. We note that all models had similar disagreements for *ρ* < 0.5 cm which is a known limitation of diffusion theory^13^.

**Figure 5.**
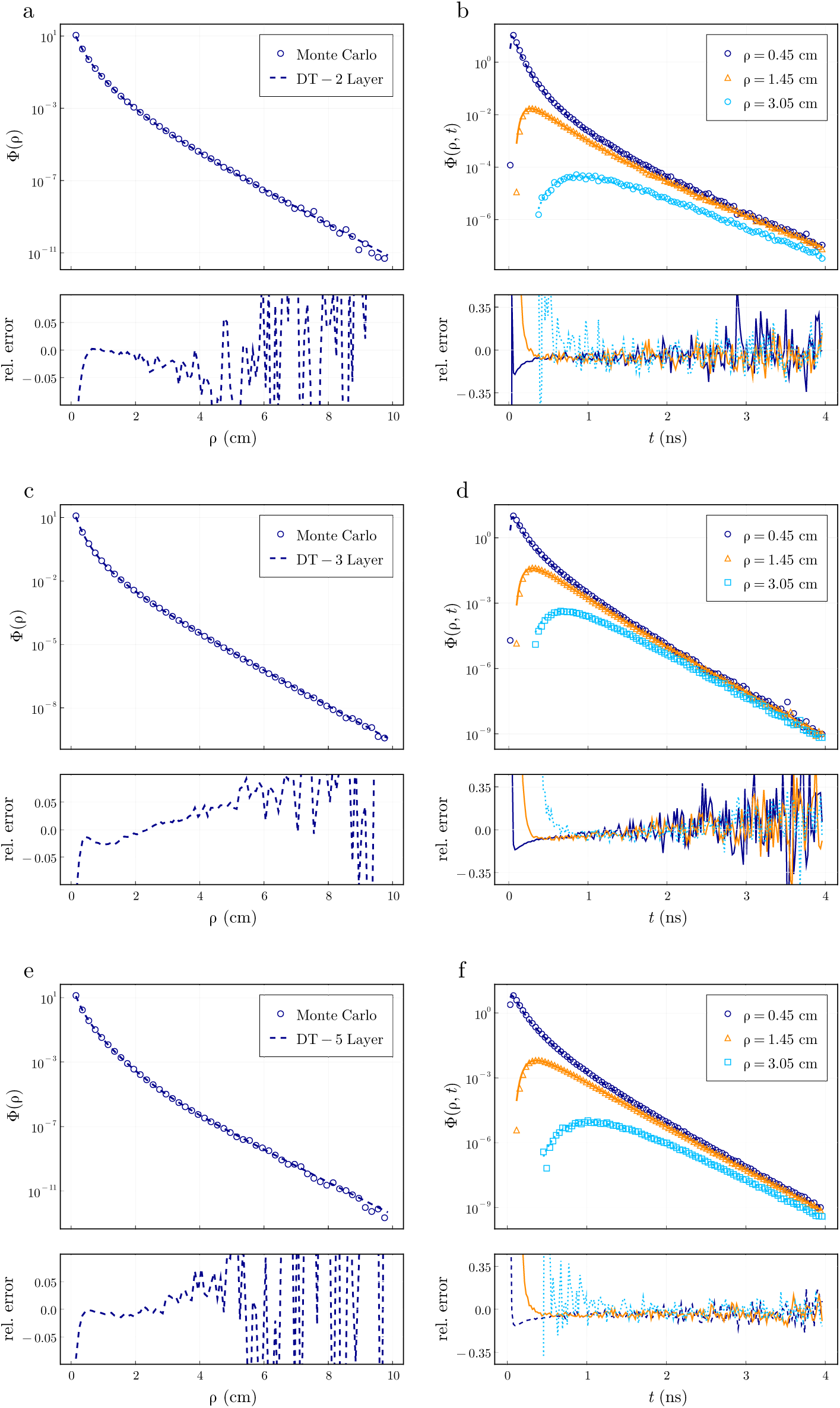
Comparison of the (left column) steady-state and (right column) time-domain fluence using diffusion theory (lines) simulated using equations (2) and (3) and the Monte Carlo method (symbols) for the tissue geometries representing a (top row) 2-layer, (middle row) 3-layer muscle, and (bottom row) 5-layer brain tissue models. The relative error between the Monte Carlo results and diffusion model are shown in the plots below. The diffusion approximation displayed relative errors less than 0.1 over a large domain of arguments suggesting it could be used in a variety of diverse tissue geometries.

In the right column of Fig. 5, we show the time-domain fluence for *ρ* = 0.45, 1.45, 3.05 cm simulated using equation (3) and compare to Monte Carlo results. The 2 layer model is well approximated by diffusion theory in the time-domain for each value of *ρ* given enough scattering events illustrated by the uniform agreement across a wide range of time values. As in the spatial domain, the time domain results for the 3-layered model do not asymptotically converge to Monte Carlo simulations due to the small top layer thickness. The agreement is not uniform for each value of *ρ* as shorter distances are better approximated until much later arrival times. This is in contrast to the 5-layer model where the errors are relatively flat at all times and distances. This could be attributed to only presenting results in the time-domain for *ρ* < 3.05 cm, whereas the effect of the low scattering CSF layer is more significant for *ρ* > 6 cm. We note that the precise choice of 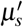 in the CSF layer for the distances and times shown do not significantly affect the time-domain simulations compared to the steady-state results. Additionally, for all comparisons against the Monte Carlo method, the finite discretization of *t, ρ* and *z* will have some effect on the resulting relative errors.

### Computational Time

In Table 2, we show the amount of time in microseconds to compute the steady-state fluence in the top layer, Φ_1_(*ρ, z* = 0), for a given number of terms *n* considered in the sum in equation (2) for 2, 4, 8 and 16 layers. Different values of optical properties do not significantly affect the computation time, which is instead dependent on the number of roots *n* used in the sum. This relation is not linear as it is faster to compute the higher order roots with asymptotic expansion for large arguments in *J*_0_(*s*_*n*_*ρ*). However, the calculation of *J*_0_(*s*_*n*_*ρ*) accounts for nearly 40 % of the run time and the computation of *G*_1_(*s*_*n*_, *z*) takes most of the remaining time. For realistic applications where less than 1,000 roots are needed, the fluence can be calculated in less than ≈ 100 *µ*s for up to 8 layers.

**Table 2.** Number of microseconds to compute steady-state fluence where *n* is number of roots in equation (2).

| Layers | $n = 100$ | $n = 500$ | $n = 1,000$ | $n = 5,000$ |
| --- | --- | --- | --- | --- |
| 2 | 8 | 33 | 64 | 268 |
| 4 | 9 | 38 | 75 | 341 |
| 8 | 13 | 60 | 118 | 469 |
| 16 | 19 | 93 | 184 | 850 |

The time-domain routine must compute the steady-state routine for *N* (≈ 12 − 24) complex absorption values. This procedure lends itself well to parallelism as each computation is independent and can be used at each time point needed. Therefore, performance is limited by the runtime listed in Table 2, however using a complex absorption term increases the runtime by 2.5x. In Table 3, we show the runtime in microseconds for the time-domain fluence as a function of the Laplace space evaluations *N* for 2 and 4 layered media considering *n* = 600 roots in equation (2) and 1024 time points. We note that the time values do not have to be linearly spaced as when using the Fast Fourier Transform.

**Table 3.** Number of microseconds to compute the time-domain fluence where *N* is the number of Laplace space evaluations.

| Layers | $N = 8$ | $N = 16$ | $N = 32$ | $N = 64$ |
| --- | --- | --- | --- | --- |
| 2 | 262 | 490 | 930 | 1,860 |
| 4 | 298 | 558 | 1,090 | 2,130 |

If a dynamic range of 3 orders of magnitude is needed, the fluence can be simulated in less than 300 *µ*s. The performance in the time-domain is highly dependent on the CPU used and its multi-core performance. When using an Intel CPU 8700k with 6 cores and 12 threads the runtimes can be decreased by 30% compared to using a MacBook Pro M1 as shown in Tables 2 and 3 using a MacBook Pro M1. The number of Laplace evaluations used should be in multiples of the available number of threads. We note that the load times for multi-threaded applications represent a significant portion of the total runtime. For a low number of Laplace evaluations (<16) these computational times can be reached within 2x using a single core. The advantage of these procedures is that rapid simulation can be performed on a personal laptop while allowing for time-domain runtimes to be significantly reduced with higher end CPUs.

## Discussion and Conclusion

Limitations of homogeneous tissue-models to describe light transport in layered biological media have been discussed previously^4,43^. Although analytical models that incorporate heterogeneous optical properties are becoming more frequent^28,44,45^, their use, particularly in inverse calculations, is limited by their numerical accuracy and efficiency^27^. Therefore, homogeneous models are typically used given their simplicity and efficiency in solving inverse problems that require 100-1,000 evaluations of the forward model to reach convergence. Several solutions for photon diffusion in layered media have been reported, but present technical difficulties for numerical computation. We have investigated a previously developed model^21,35^ that has received wide interest^28,44–46^. However, the model relies on numerical inverse transforms for obtaining the photon fluence for both steady-state and time-domain simulations which limits the numerical accuracy and speed. For example, the computation of the steady-state fluence requires the inversion of a Bessel-type 1-D inverse transform (equation 2) over the *k*^*th*^ root of the zeroth order Bessel function *J*_0_. The discrete version has several advantages compared to using Gaussian integration^19,22^ as the roots can be precomputed to improve the overall speed of the routine and can be accurately computed over a wide range of input arguments using a variable number of roots. This is important because the convergence of equation (2) is highly dependent on the model inputs (Fig. 2), where different optical properties and tissue geometries require a different number of roots to be used. However, these expressions require numerical integration over hyperbolic functions that can numerically overflow for large input arguments or at large roots of *J*_0_.

In this work, we provide numerically stable expressions for the Green’s functions in terms of exponentially decaying functions, which facilitates accurate computation for large input arguments (e.g., scattering, layer thickness, spatial frequency) over any root of *J*_0_ without approximations or loss of generality that are usually required to numerically compute equation (2)^28,29^. As shown in Fig. 2, the accuracy and speed of computed solutions is determined by the number *n* of roots used in the sum in equation (2) which is dependent on 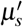, *a*, and *z*. In practice, the values used for the cylindrical radius of the tissue-model *a* (Fig 2d) should be kept as small as possible to increase convergence but should be large enough to accurately represent lateral boundary conditions. Although the total number of *n* largely dictates the speed and accuracy of the routine, the algorithm is limited to simulating equation (2) with absolute errors up to the machine precision in the calculation. This is largely due to the finite precision used in the calculation of *J*_0_(*s*_*n*_*ρ*) in equation (2) which is limited to absolute tolerances approaching the machine precision. For higher precision calculations, it is important to calculate the roots of *J*_0_ in the desired precision to simulate fluence values down to the machine precision. For experimental measurements with background noise, fluence values below the epsilon value of double precision (*ε* ≈ 10^−16^) are rarely needed. Additionally, computing *J*_0_ accounts for the majority of the routine’s runtime especially when the fluence is required at multiple spatial locations, as is the case in many tomography^47^ or functional imaging^48^ applications. New numerical routines for the computation of *J*_0_ were developed that decrease computational time by at least 3x^49^ compared to using standard routines^50^. An advantage of the routine presented is that computing the fluence at 10 arbitrarily specified spatial locations takes only 3x longer than the times reported in Table 2. Although it can be difficult to directly compare computational times of different routines, as they depend highly on the computational resources and effort put into them, we were able to simulate the steady-state fluence 500-1,000x faster than previously reported^19,46^. We note that these times are achieved on a personal laptop using a single core.

Computation of the time-domain signal requires an additional inverse time transform which is usually performed with the Fourier Transform^28^. Here, we have used the inverse Laplace transform^33,34^ for faster and more accurate reconstructions of the time-domain signal. We have found that 12-24 terms in the Laplace integral are needed in equation 3) to reconstruct the time-domain signal with dynamic ranges of 3-6 orders of magnitude, which is the range of current experimental systems^8,39^. Due to the decreased number of evaluations needed in the inverse time transform, the computational times for time-domain simulations are 1,000-10,000x faster than what is usually reported depending on the number of layers considered and accuracy required^21,24,31^. Most of the performance gain can be attributed to utilizing the faster converging Laplace transform instead of the Fourier transform^33^ while other improvements come from other numerical optimizations for the steady-state calculation and threaded parallelism as further discussed in Appendix A in the Supplementary material. The Laplace transform can also evaluate the time-domain fluence up to absolute errors approaching the machine precision as shown in Fig. (3). The number of terms needed in the Laplace transform for adequate convergence will depend highly on the contour size Λ = *t*_2_/*t*_1_ which is recommended to be kept as small as possible for faster reconstructions.

A primary limitation of the layered solutions presented here is that a large amount (500-5,000x) of terms are required in the computation of equation (2) when *z* = 0 which is required for reflectance calculations. As seen in Fig. 2, increasing the top layer scattering coefficient will significantly increase the number of terms required in equation (2), while the convergence is mostly independent of deeper layer optical properties. This can be explained by the slow convergence of the particular solution of the Green’s function when 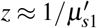. When *z* is farther away from the source depth *z*_0_ as seen in Fig. 2c, only a few terms are needed. However, if we approximate that *z* ≈ *z*_0_, it becomes possible to sum the particular solution of the Green’s function exactly which improves convergence significantly. We present detailed derivations of this approximation in Appendix A in the Supplementary material and show that such an approximation when 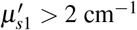 and *z* = 0 can simulate the fluence with relative errors down to 10^−14^ (Supplementary Fig. S3), which is as accurate as the exact forms in double precision arithmetic due to floating point errors. Therefore, it is highly recommended to use such an approximation in double precision arithmetic which can decrease computational times by 2-3 orders of magnitude depending on the input 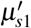, allowing for computation of the steady-state fluence in less than a microsecond ((Supplementary Fig. S4). This approximate form also allows for very accurate simulation for large scattering coefficients where it is difficult for the exact expressions to converge due to the slow exponential decay of the sum.

Finally, in addition to testing the numerical accuracy and efficiency, we have tested the physical approximation of diffusion theory compared to the Monte Carlo method by using three previously reported layered tissue models that approximate several different organ systems^22,30,40^. We note that solutions to the RTE in layered media have been presented which are more accurate than the diffusion approximation but, like Monte Carlo methods, come at increased computational cost^41^. Additionally, the situations presented in Fig 5 represent the simplest forms of heterogeneous media consisting of layered slabs which may be a rather crude approximation of complex biological media. Though, the use of such a simple approximation has been shown to provide similar accuracy to more realistic tissue geometries in a brain model using atlas based meshes^3^. The primary disadvantages of diffusion theory are the inability to correctly predict photon fluence for short time scales and source-detector separations. A recent report also indicated that the diffusion approximation could increase inaccuracies far away from the source in layered models where layer thicknesses are small compared to the mean scattering length^41^. We found that for a top layer thickness of 1.2 mm, these predictions were in agreement with the results reported in Fig. 5, but we also find that such errors were only significant at distances of *ρ* > 5 cm for steady-state calculations. These errors were not apparent in the time-domain for *ρ* = 0.45, 1.45, 3.05 cm when *t* < 4 ns (Fig. 5). We also find that our solutions from diffusion theory agree well with Monte Carlo simulations for a 5-layer model of the brain even when considering a thin CSF layer of low scattering (Fig. 5).

In conclusion, we have developed and verified an open-source, easy-to-use numerical algorithm to accurately and efficiently compute solutions of the diffusion equation in layered media. The absolute errors of the routine can be made arbitrarily accurate and can simulate both the steady-state and time-domain fluence 3 to 4 orders of magnitude faster than previously reported. Therefore, the routine can be used in inverse procedures to recover optical properties of measured data in real-time (1-10 Hz). It can also be employed for rapid generation of the intensity profile in layered media at multiple spatial locations and varying optical properties, as required in tomography and functional imaging applications. These solutions are also easily amendable to solve the correlation diffusion equation in layered media. An additional advantage of the routine is that the computational time marginally increases with the addition of a new layer, as a 4-layered medium can be computed within 10% of the time to compute 2-layers. This could allow for more accurate simulations in highly layered media such as the brain at little cost to total runtimes. Additionally, we showed good agreement between diffusion theory and Monte Carlo simulations in three separate tissue geometries of clinical interest.

## Code Availability

All software used in this manuscript are freely available online with documentation at https://github.com/heltonmc/LightPropagation.jl. The code for the numerical inversion of the Laplace Transform is also available at https://github.com/heltonmc/Laplace.jl along with several other algorithms not shown in this manuscript to invert the Laplace transform. The code for calculating Bessel’s functions is available at https://github.com/heltonmc/Bessels.jl

## Acknowledgements

The authors would like to thank Dr. Robert H. Wilson for helpful comments on earlier drafts of this manuscript. The authors are greatly appreciative of Dr. André Liemert for helpful discussions on the theoretical solutions and to Dr. Carole Hayakawa for help in setting up the Monte Carlo simulations used in this work.

## Author contributions statement

M.H. developed the numerical code and theoretical solutions, M.H. and S.Z. conducted Monte Carlo analysis, M.H. and S.Z. analysed all data, M.H., K.V. and M-A.M. designed the studies, all authors reviewed and contributed to writing the manuscript. K.V. and M-A.M. supervised this work.

