## Supplementary Material for "Light diffusion in layered media: A numerical study in the spatial and time-domains"

### ABSTRACT

Accurate and efficient forward models of photon migration in heterogeneous geometries are important for many applications of light in medicine because many biological tissues exhibit a layered structure, with each layer having independent optical properties and thickness. Unfortunately, closed form analytical solutions are not readily available for layered tissue-models, and often are modeled using computationally expensive numerical techniques or theoretical approximations that limit accuracy and real-time analysis. Here, we develop an open-source accurate, efficient, and stable numerical routine to solve the diffusion equation in the steady-state and time-domain for a layered cylinder tissue model with an arbitrary number of layers and specified thickness and optical coefficients. We show that the steady-state ($< 0.1$ ms) and time-domain ($< 0.5$ ms) fluence (for an 8-layer medium) can be calculated with absolute numerical errors approaching machine precision. The numerical implementation increased computation speed by 3 to 4 orders of magnitude compared to previously reported theoretical solutions in layered media. We verify our solutions asymptotically to homogeneous tissue geometries using closed form analytical solutions to assess convergence and numerical accuracy. Approximate solutions to compute the reflected intensity are presented which can decrease the computation time by an additional 2-3 orders of magnitude. We also compare our solutions for 2, 3, and 5 layered media to gold-standard Monte Carlo simulations in layered tissue models of high interest in biomedical optics (e.g. skin/fat/muscle and brain). The presented routine could enable more robust real-time data analysis tools in heterogeneous tissues that are important in many clinical applications such as functional brain imaging and diffuse optical spectroscopy.

### Supplementary Material

### Appendix A: Layered solutions in Steady-State

We use the integral transform approach[1] to solve the diffusion equation for a N-layered cylindrical model as shown in Fig. S1. A collimated source-beam is approximated by an isotropic point source located at a distance of $z_0 = 1/\mu'_{s1}$ from the location of incidence of the beam and boundary with $\mu'_{s1}$ representing the reduced scattering coefficient in the first layer.

The steady-state diffusion equation can be given by

$$D\nabla^2\Phi(\vec{r}) - \mu_a\Phi(\vec{r}) = -S(\vec{r}) \tag{A.1}$$

where $\Phi$, $D = 1/(3\mu'_s)$, and $\mu_a$ denote the fluence rate, the diffusion coefficient, and the absorption coefficient, respectively[1].

The source function $S(\vec{r})$ can be expressed as a Dirac delta function in cylindrical coordinates. Eq. A.1 can then be rewritten in cylindrical coordinates as

$$\frac{\partial^2}{\partial\rho^2}\Phi + \frac{1}{\rho}\frac{\partial}{\partial\rho}\Phi + \frac{1}{\rho^2}\frac{\partial^2}{\partial\phi^2}\Phi + \frac{\partial^2}{\partial z^2}\Phi - \frac{\mu_a}{D}\Phi = -\frac{1}{D\rho}\delta(\rho-\rho_0)\delta(\phi-\phi_0)\delta(z-z_0) \tag{A.2}$$

with the abbreviation $\Phi = \Phi(\rho,\phi,z)$. The derivative with respect to $\phi$ in Eq. A.2 can be eliminated using a cosine transform and then reduced to an ordinary differential equation by using the finite Hankel transform of $m^{th}$ order[2]. These two transforms can be expressed together by[1]

$$\Phi(s_n,\phi,m) = \int_0^{2\pi}\int_0^{a'} \rho\Phi(\rho,\phi')J_m(s_n\rho)\cos(m(\phi-\phi'))\,d\rho\,d\phi' \tag{A.3}$$

using an extrapolated boundary condition at $\rho = a'$ for the upper limit of the integral transform with $a' = a + z_{b1}$ where $a$ is the radius of the cylinder. The extrapolation length can be calculated with $z_{bk} = 2AD_k$ where $A$ is proportional to the fraction

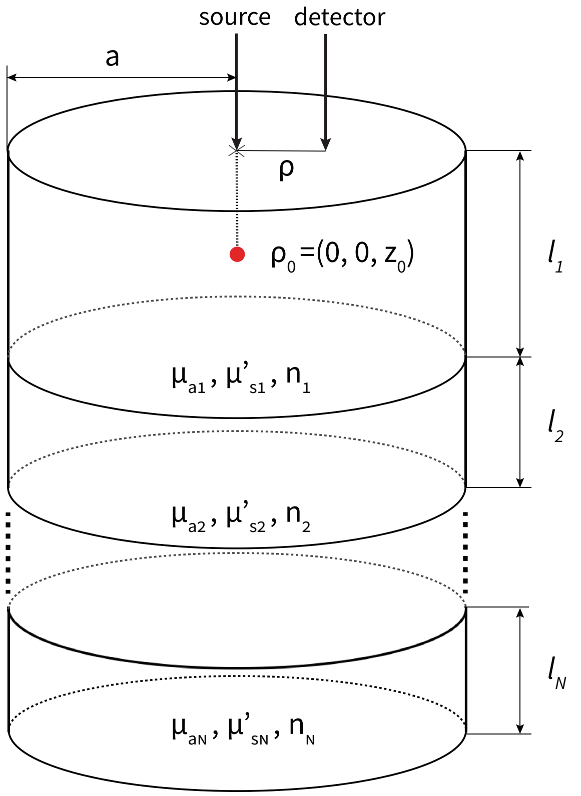


**Supplementary Figure 1.** Schematic of the $N-$layered turbid medium with a source located onto the center of the cylinder top.

of photons that are internally reflected at the boundary[3]. The subscript $k$ signifies the $k^{th}$ layer of the cylinder with distinct absorption $\mu_{ak}$ and scattering $\mu'_{sk}$ in each layer. Applying Eq. A.3 to Eq. A.2 yields an ordinary differential equation

$$\frac{\partial^2}{\partial z^2}\Phi - (\frac{\mu_a}{D} + s_n^2)\Phi = -\frac{1}{D}J_m(s_n\rho_0)\cos(m(\phi-\phi_0))\delta(z-z_0) \tag{A.4}$$

for $\Phi = \Phi(s_n, \phi, m, z)$ after applying a finite Hankel transform relation[2]. $J_m$ is the Bessel function of first kind and order $m$ and each $s_n$ is determined from the roots of $J_m$ such that $J_m(a's_n) = 0, \quad n = 1, 2, ...,$.

A Green's function approach is used solve Eq. A.4 for the fluence in a specific layer assuming that the isotropic source (from an incident pencil beam) is located within the first layer ($0 \leq z_0 < l_1$). We then seek separate Green's functions in each layer where the solution for the first layer $G_1(s_n, z)$ is composed of a homogenous and particular part while the solutions for the remaining $k$ layers have only a homogeneous solution. Therefore, the Green's function $G_1(s_n, z)$ in the top layer becomes

$$\frac{\partial^2}{\partial z^2}G_1(s_n,z) - \alpha^2 G_1(s_n,z) = -\frac{1}{D_1}\delta(z-z_0) \tag{A.5}$$

when $z$ is within the first layer $0 \leq z < l_1$. For the $N^{th}$ layer, $G_N(s_n, z)$ becomes

$$\frac{\partial^2}{\partial z^2}G_N(s_n,z) - \alpha^2 G_N(s_n,z) = 0 \tag{A.6}$$

when $z$ is located in the bottom layer $\sum_{k=1}^{N-1} l_k \leq z < \sum_{k=1}^{N} l_k$. These equations can be used to reduce Eq. A.4 to

$$\Phi_k(s_n,\phi,m,z) = G_k(s_n,z)J_m(s_n\rho_0)\cos(m(\phi-\phi_0)) \tag{A.7}$$

We use the boundary conditions previously described[1] but provide solutions for $G_1(s_n, z)$ and $G_N(s_n, z)$ purely in terms of exponentially decaying functions. This form of expression provides stability in numerical calculations and is in contrast to previously derived expressions[1] which contain hyperbolic trignometric functions that can easily produce overflow-errors. We also note that the solutions we derive are exact and do not require any approximations for calculation, as is required by previous reports[4–6]. Expanding hyperbolic functions as exponentials also allows for simplifying several other terms that serve to reduce the computational time.

The Green's function $G_1$ in the first layer $(0 \leq z < l_1)$ is given by

$$G_1(s_n, z) = \frac{e^{-\alpha_1|z-z_0|} - e^{-\alpha_1(z+z_0+2z_{b1})}}{2D_1\alpha_1} + \frac{e^{\alpha_1(z+z_0-2l_1)}(1-e^{-2\alpha_1(z_0+z_{b1})})(1-e^{-2\alpha_1(z+z_{b1})})}{2D_1\alpha_1} \times \frac{D_1\alpha_1 n_1^2\beta_3 - D_2\alpha_2 n_2^2\gamma_3}{D_1\alpha_1 n_1^2\beta_3(1+e^{-2\alpha_1(l_1+z_{b1})}) + D_2\alpha_2 n_2^2\gamma_3(1-e^{-2\alpha_1(l_1+z_{b1})})} \tag{A.8}$$

where $\alpha_k = \sqrt{\mu_{ak}/D_k + s_n^2}$. In general, the quantities $\beta_3$ and $\gamma_3$ are obtained by downward recurrence relations with start values

$$\begin{aligned} \beta_N &= D_{N-1}\alpha_{N-1}n_{n-1}^2(1+e^{-2\alpha_{N-1}l_{N-1}})(1-e^{-2\alpha_N(l_N+z_{b_2})}) \\ &\quad + D_N\alpha_N n_n^2(1-e^{-2\alpha_{N-1}l_{N-1}})(1+e^{-2\alpha_N(l_N+z_{b_2})}) \\ \gamma_N &= D_{N-1}\alpha_{N-1}n_{n-1}^2(1-e^{-2\alpha_{N-1}l_{N-1}})(1-e^{-2\alpha_N(l_N+z_{b_2})}) \\ &\quad + D_N\alpha_N n_n^2(1+e^{-2\alpha_{N-1}l_{N-1}})(1+e^{-2\alpha_N(l_N+z_{b_2})}) \end{aligned} \tag{A.9}$$

with the downward recurrence given by

$$\begin{aligned} \beta_{k-1} &= D_{k-2}\alpha_{k-2}n_{k-2}^2(1+e^{-2\alpha_{k-2}l_{k-2}})\beta_k + D_{k-1}\alpha_{k-1}n_{k-1}^2(1-e^{-2\alpha_{k-2}l_{k-2}})\gamma_k \\ \gamma_{k-1} &= D_{k-2}\alpha_{k-2}n_{k-2}^2(1-e^{-2\alpha_{k-2}l_{k-2}})\beta_k + D_{k-1}\alpha_{k-1}n_{k-1}^2(1+e^{-2\alpha_{k-2}l_{k-2}})\gamma_k \end{aligned} \tag{A.10}$$

We note that we seek just the terms $\beta_3$ and $\gamma_3$ which must be determined recursively if the total number of layers $N$ is larger than 3. In that case, Eq. A.9 is used to generate starting values in the recurrence relation, then Eq. A.10 is used recursively until $\beta_3$ and $\gamma_3$ are obtained. If $N = 2$, $\beta_3 = 1 - e^{-2\alpha_2(l_2+z_{b2})}$ and $\gamma_3 = 1 + e^{-2\alpha_2(l_2+z_{b2})}$. For $N = 3$, only Eq. (A.9) is needed to calculate $\beta_3$ and $\gamma_3$.

The solution to Eq. A.11 for $G_N$ is

$$G_N(s_n, z) = \frac{n_N^2 2^{N-2}\prod_{i=2}^{N-1}(D_i\alpha_i n_i^2)\exp(\alpha_1(z_0 - l_1) + \alpha_N(L_N + z_{bN} - z) - \xi_2)}{D_1\alpha_1 n_1^2\beta_3(1+e^{-2\alpha_1(l_1+z_{b1})}) + D_2\alpha_2 n_2^2\gamma_3(1-e^{-2\alpha_1(l_1+z_{b1})})} \tag{A.11}$$

The quantity $\xi_2$ is computed with start values $\xi_N = \alpha_N(l_N + z_{b2})$ and with downward recurrence relations $\xi_{k-1} = \xi_k + \alpha_{k-1}l_{k-1}$ such that for $N = 2$, $\xi_2 = \alpha_2(l_2 + z_{b2})$ and for $N = 3$, $\xi_2 = \alpha_2 l_2 + \alpha_3(l_3 + z_{b2})$ and so on. We note that our routine has unrolled these relations completely for $N = 4$ to improve performance.

For solutions in real-space, $\Phi(\rho, z)$ we apply the inverse relation of Eq. A.3 to Eq. A.7 such that the fluence in real space can be written as[1]

$$\Phi_k(\rho, z) = \frac{1}{\pi a'^2}\sum_{n=1}^{\infty} G_k(s_n, z)J_0(s_n\rho)J_1^{-2}(a's_n) \tag{A.12}$$

for the special case of a point source that is incident at the center top of the cylinder.

### Computing the real-space fluence

We note a few points that help to efficiently compute Eq. A.12. First, we determine $s_n$ such that $J_0(a's_n) = 0, \quad n = 1, 2, ...,$. Calculation of these roots at runtime significantly impacts performance. Instead $n$ roots of $J_0(x)$ are precomputed and the relation $s_n = j_{0,n}/a'$ where $j_{0,n}$ is the $n^{th}$ root of $J_0(x)$ is calculated at runtime. It is then also possible to remove the runtime computation of $J_1^{-2}(a's_n)$ because $a's_n = j_{0,n}$ such that $J_1^{-2}(j_{0,n})$ is constant and therefore can be precomputed. For results shown in this manuscript, we computed $j_{0,n}$ for $n = 1...1,000,0000$ in octuple precision which are loaded along with $J_1^{-2}(j_{0,n})$ to be used in every simulation. We have observed that simply computing the Bessel functions account for over 50% of code execution and removing a function call to $J_1(x)$ substantially reduced the total computation time. For the calculation of $J_0(x)$, we developed an optimized routine[7] that increased computation speed by 3x relative to the common Amos' Fortran routines[8]. This optimization is crucial for fast calculation of the fluence at multiple spatial locations. Loss of numerical precision in calculation of $J_0(s_n\rho)$ limits the accuracy of solutions typically to errors on the order of the machine precision. Though, we note that our routine can be run in arbitrary precision at the cost of increased computational time. Lastly, it may also be advantageous if computing the fluence at fixed spatial locations to precompute $J_0(\rho j_{0,n}/a')$ for several values of $\rho$ by approximating that $a' \approx a$ if the cylinder radius is large. This would eliminate any calls within the loop to special function libraries which would significantly improve performance while allowing for calculation at varying optical properties and layer thicknesses as required in inverse problems.

#### Approximate solutions for $\mathbf{z \approx z_0}$

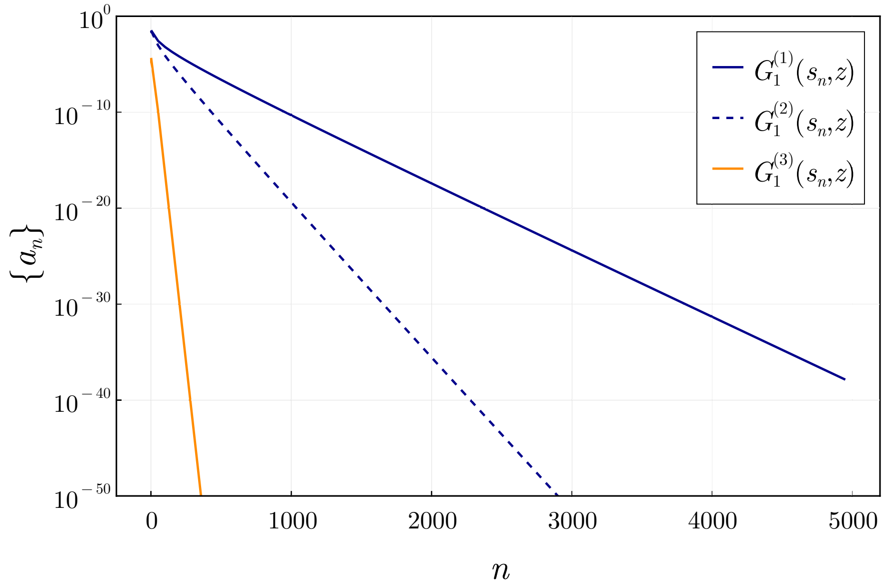


**Supplementary Figure 2.** We show the convergence of three separate terms in Eq. A.8 by showing the value of the $n^{th}$ term in the sequence when summing over $j_{0,n}$. Each term converges at a much different rate with the overall convergence being highly dependent on $\mu'_{s1}$ when $z = 0$.

A limitation of computing $\Phi_1(\rho, z)$ using $G_1(s_n, z)$ in Eq. A.8 is the slow convergence when $z \approx z_0$. This is of particular importance for reflectance measurements when we must compute $\Phi_1(\rho, z)$ when $z = 0$. Unfortunately, when $z = 0$ and $\mu'_{s1}$ is large, the routine is limited by slow convergence requiring thousands of terms in Eq. A.12. The convergence of Eq. A.12 for $\Phi_1(\rho, z)$ can be analyzed by separating Eq. A.8 into three terms

$$G_1^{(1)}(s_n, z) = \frac{e^{-\alpha_1|z-z_0|}}{2D_1\alpha_1} \tag{A.13}$$

$$G_1^{(2)}(s_n, z) = \frac{-e^{-\alpha_1(z+z_0+2z_{b1})}}{2D_1\alpha_1} \tag{A.14}$$

$$\begin{aligned} G_1^{(3)}(s_n, z) = &\frac{e^{\alpha_1(z+z_0-2l_1)}(1-e^{-2\alpha_1(z_0+z_{b1})})(1-e^{-2\alpha_1(z+z_{b1})})}{2D_1\alpha_1} \\ &\times \frac{D_1\alpha_1 n_1^2\beta_3 - D_2\alpha_2 n_2^2\gamma_3}{D_1\alpha_1 n_1^2\beta_3(1+e^{-2\alpha_1(l_1+z_{b1})}) + D_2\alpha_2 n_2^2\gamma_3(1-e^{-2\alpha_1(l_1+z_{b1})})} \end{aligned} \tag{A.15}$$

where $G_1(s_n, z) = G_1^{(1)}(s_n, z) + G_1^{(2)}(s_n, z) + G_1^{(3)}(s_n, z)$. In Fig. S2, we show the value of the three terms $a_n$ as we iterate over the $n^{th}$ root of $j_{0,n}$. Here, we show a specific example when $\mu'_{s1} = 20$ cm$^{-1}$, $\mu'_{s2} = 15$ cm$^{-1}$, $\mu_{a1} = 0.1$ cm$^{-1}$, $\mu_{a2} = 0.2$ cm$^{-1}$, $a = 10$ cm, $l_1 = 0.5$ cm, and $l_2 = 5$ cm. We can see that $G_1^{(3)}(s_n, z)$ is rapidly decaying and just a few terms are needed for accurate computation. It should also be noted that this term is the only term affected by the optical properties of deeper layers $k \geq 2$ and therefore the dominate convergence of Eq. A.8 is only affected by the optical properties of the first layer. However, because the three terms decay at such a different rate, for the highest numerical accuracy it is recommended to sum these terms separately and combine them once the infinite sum is terminated. As we can see, the convergence is dominated by the slow decay of $G_1^{(1)}(s_n, z)$ when $z = 0$. This convergence becomes slower as $\mu'_{s1}$ becomes larger.

However, when $z \approx z_0$ we can sum the particular solution $G_1^{(1)}(s_n, z)$ over $n$ exactly in closed form yielding the infinite space Green's function caused by an isotropic point source. This allows us to write the fluence $\Phi_1(z \approx z_0)$ as

$$\Phi_1(\rho) = \frac{e^{-\kappa_1 |r|}}{4\pi D_1 |r|} + \Phi_1^{(h)}(\rho) \tag{A.16}$$

where $r = \sqrt{\rho^2 + (z - z_0)^2}$ and $\kappa_1 = \sqrt{\mu_{a1}/D_1}$. The first term in Eq. A.16 is the infinite space Green's function. The homogenous part $\Phi_1^{(h)}(\rho)$ can be computed with Eq. A.12 and $G_1^{(h)}(s_n, z)$ computed with $G_1(s_n, z) = G_1^{(2)}(s_n, z) + G_1^{(3)}(s_n, z)$. We note that this solution is only approximate if we want to consider the fluence on the boundary $z = 0$, but to calculate the fluence inside the first layer when $z = z_0$ it represents the exact solution. This significantly reduces the number of terms needed in Eq. A.12, however $G_1^{(2)}(s_n, z)$ now limits the overall convergence rate which also decays at a slow rate when $z = 0$ and for increasing $\mu'_{s1}$. However, a similar approximation can be made which allows for the exact summation of $G_1^{(1)}(s_n, z) + G_1^{(2)}(s_n, z)$ in closed form which represents the steady-state solution for a semi-infinite medium. Therefore, it is appropriate to rewrite the fluence as

$$\Phi_1(\rho, z = 0) \approx \Phi^{SI}(\rho, z) + \frac{1}{\pi a'^2} \sum_{n=1}^{\infty} G_1^{(3)}(s_n, z) J_0(s_n \rho) J_1^{-2}(a' s_n) \tag{A.17}$$

where $\Phi^{SI}(\rho, z)$ is the steady-state fluence in a semi-infinite medium (Equation 3 given in Kienle and Patterson[9]). We test this approximation in Fig. S3 as a function of $\mu'_{s1}$ and $\rho$ by summing $G_1^{(1)}(s_n, z) + G_1^{(2)}(s_n, z)$ for $n = 50{,}000$ in octuple precision using Eq. A.12 and comparing the absolute and relative errors to the closed form semi-infinite Green's function. In Fig. S4, we compare the approximation to the exact solution for two tissue models: (a) $\mu'_{s1}$ = 10 cm$^{-1}$, $\mu'_{s2}$ = 13 cm$^{-1}$, $\mu_{a1}$ = 0.1 cm$^{-1}$, and $\mu_{a2}$ = 0.2 cm$^{-1}$ and (b) $\mu'_{s1}$ = 60 cm$^{-1}$, $\mu'_{s2}$ = 40 cm$^{-1}$, $\mu_{a1}$ = 0.01 cm$^{-1}$, and $\mu_{a2}$ = 0.08 cm$^{-1}$. We use the same layer thicknesses, $l_1 = 1$ and $l_2 = 5$ cm with a cylinder radius of 20 cm.

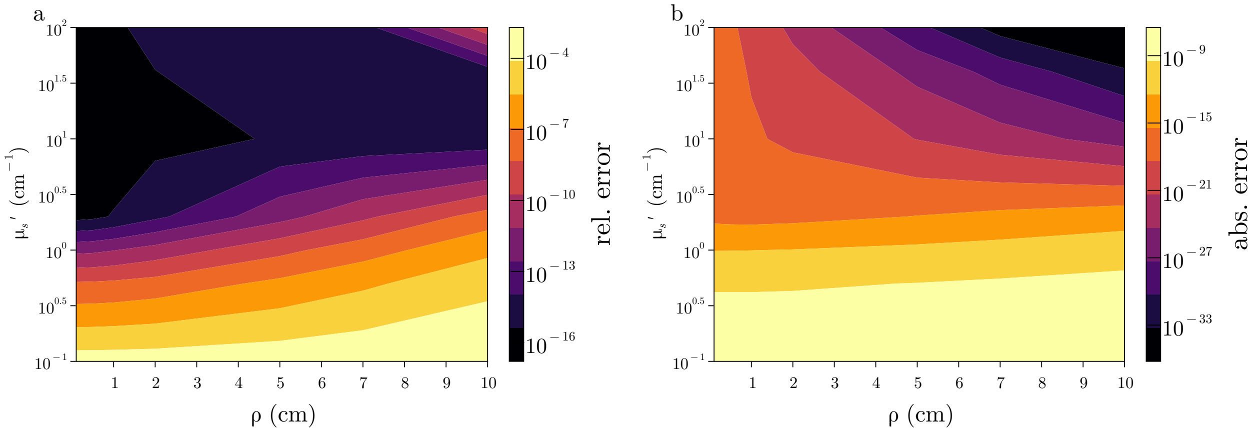


**Supplementary Figure 3.** The (left) relative and (right) absolute error between the closed-form semi-infinite Green's function and $G_1^{(1)}(s_n, z) + G_1^{(2)}(s_n, z)$ when summed over 50,000 terms using Eq. A.12. This approximation also gives absolute errors below the machine precision in double precision calculations when $\mu'_{s1} > 2$ cm$^{-1}$.

Theoretically, this solution becomes more accurate when $z = 0$ as $\mu'_{s1} \to \infty$. We find this approximation to be accurate for relative errors greater than $10^{-14}$ for $\mu'_{s1} > 2$ cm$^{-1}$ which is within the relative errors provided by the exact forms when double precision arithmetic is used. Even for very low $\mu'_{s1} = 0.1$ cm$^{-1}$, this approximation can give at least 3 digits of accuracy. The advantage of this approach is that just 150 roots were used in the approximate form when $\mu'_s = (10, 13)$ cm$^{-1}$ where 3,500 roots were needed in the exact form. When $\mu'_s = (60, 40)$ cm$^{-1}$ the approximation only needed 200 roots at all values of $\rho$ compared to 20,000 roots required in the exact form. However, if only a couple digits of accuracy are needed the approximate form can give reasonable convergence in less than 50 roots even for very large scattering coefficients. This results in computational times being 2-3 orders of magnitude faster allowing for computation in less than one microsecond for a wide range of optical properties. Therefore, it is highly recommended to use such an approximation for $\mu'_{s1} > 2$ cm$^{-1}$.

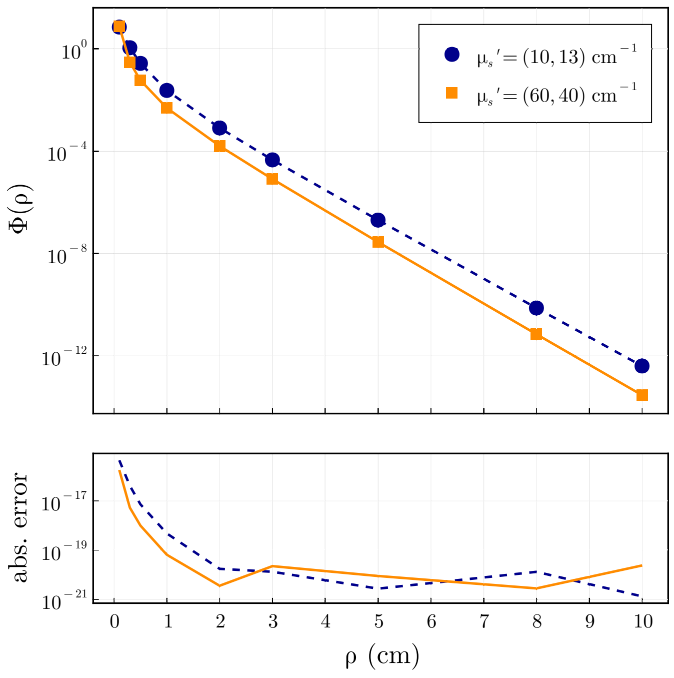


**Supplementary Figure 4.** Shows the steady-state fluence calculated with the semi-infinite space Green's function (markers) compared to summing $G_1^{(1)}(s_n,z) + G_1^{(2)}(s_n,z)$ exactly using Eq. A.12 (lines) for two different tissue geometries. The absolute error is shown in the below plot displaying that these solutions give absolute errors below the machine precision in double precision calculations. The solutions were computed in octuple precision for accurate comparison.

### Appendix B: Layered solutions in Time-Domain

Given a sinusoidally modulated source at frequency $f$, the real and imaginary parts of the fluence $\Phi(\rho,\omega)$ can be calculated using the same formula for $\Phi(\rho)$ and adding a complex absorption term[10]

$$\alpha_k = \sqrt{\mu_{ak}/D_k + s_n^2 + i\omega/(D_k c)} \tag{A.18}$$

where $c$ is the speed of light in the medium and $i = \sqrt{-1}$. The real and imaginary parts are used to calculate the phase angle and modulation.

For solutions in the time-domain, the real and imaginary parts of the fluence in the frequency domain must be calculated at many frequencies (400-4,000x)[10,11] and inverse Fourier transformed into the time-domain[2]. The Fourier integral is slowly converging and the number of frequency evaluations needed is highly dependent on $\rho$, $\mu_s'$ and $t$[11].

Alternately, an inverse Laplace transform can be applied to Eq. A.12[11,12] by making the substitution $i\omega \to \bar{s}$ in Eq. A.18 and numerically integrating the Bromwich complex contour integral

$$f(t) = \frac{1}{2\pi i}\int_B e^{\bar{s}t} F(\bar{s})\, d\bar{s} \tag{A.19}$$

In Eq. A.19, $B$ denotes the Bromwich path where $\bar{s}$ is a complex number along the contour. We note we use a bar $\bar{s}$ to avoid confusion with $s_n$. The corresponding solution for the time-domain fluence is then

$$\Phi_k(\rho,t) = \frac{1}{\pi a'^2}\sum_{n=1}^{\infty}\frac{1}{2\pi i}\left[\int_B e^{\bar{s}t} G_k(s_n,z,\bar{s})\, d\bar{s}\right] \times J_0(s_n\rho) J_1^{-2}(a's_n) \tag{A.20}$$

The Bromwich line can be deformed into a Hankel contour that begins and ends in the left half-plane, such that $Re\, z \to -\infty$[12]. On such a contour, the exponential in Eq. A.19 ensures that the integrand decays rapidly and renders the integral well-suited for approximation using a simple trapezoidal rule. We utilized a hyperbola countour[12] parameterized by $s(\theta) = \mu + i\mu\sinh(\theta + im\varphi)$ where $s'(\theta) = i\mu\cosh(\theta + i\varphi)$ to evaluate the time-domain solutions. For non-complex time-domain signals, application of the midpoint rule gives:

$$f(t) = \frac{h}{\pi}\left[\sum_{k=0}^{N-1} F(\bar{s}_k)\exp(\bar{s}_k t)\bar{s}'_k\right] \quad (A.21)$$

where $\bar{s}_k = \bar{s}(\theta_k)$ for $\theta_k = (k+1/2)h$ and $h$ being the uniform node spacing with N being the number of nodes along the hyperbola in the upper half-plane $Re(s) > 0$

The parameters $\mu$ and $\varphi$ as well as node spacing $h$ are obtained for computing $f(t)$ across many time values in $t \in (t_1, t_2)$ by considering a single (fixed) integration path for all time values. The time-independent parameters are expressed by:

$$\mu = \frac{4\pi\varphi - \pi^2}{A(\varphi)}\frac{N}{t_2} \quad (A.22)$$

$$A(\varphi) = \text{arcosh}\left[\frac{(\pi - 2\varphi)\Lambda + 4\varphi - \pi}{(4\varphi - \pi)\sin\varphi}\right] \quad (A.23)$$

where $\Lambda = t_2/t_1$. The uniform node spacing becomes $h = A(\varphi)/N$ with the previously optimized[11] parameter $\varphi = 1.09$.
